# Simons Collaborative Marine Atlas Project (Simons CMAP): an open-source portal to share, visualize and analyze ocean data

**DOI:** 10.1101/2021.02.16.431537

**Authors:** Mohammad D. Ashkezari, Norland R. Hagen, Michael Denholtz, Andrew Neang, Tansy C. Burns, Rhonda L. Morales, Charlotte P. Lee, Christopher N. Hill, E. Virginia Armbrust

## Abstract

Simons Collaborative Marine Atlas Project (Simons CMAP) is an open-source data portal that interconnects large, complex, and diverse public data sets currently dispersed in different formats across different Oceanography discipline-specific databases. Simons CMAP is designed to streamline the retrieval of custom subsets of data, the generation of data visualizations, and the analyses of diverse data, thus expanding the power of these potentially underutilized data sets for cross-disciplinary studies of ocean processes. We describe a unified architecture that allows numerical model outputs, satellite products, and field observations to be readily shared, mined, and integrated regardless of data set size or resolution. A current focus of Simons CMAP is integration of physical, chemical, and biological data sets essential for characterizing the biogeography of key marine microbes across ocean basins and seasonal cycles. Using a practical example, we demonstrate how our unifying data harmonization plans significantly simplifies and allows for systematic data integration across all Simons CMAP data sets.

## 1 Introduction

Oceanography is a dynamic and diverse discipline that investigates biological, chemical, and physical processes of the world’s oceans via theoretical and experimental approaches. The advent of satellite technologies and autonomous instrumentation in conjunction with ubiquitous high-performance computational resources is resulting in an explosive growth in the volume, variety and velocity of compiled oceanographic data sets. In principle, cross-disciplinary analyses of these data collections promise new scientific advances. And yet, the sheer size and complexity of available data frequently hinders the ability to fully capitalize on this collective potential. Moreover, sharing the generated data sets remains a fundamental cultural and technical challenge [1] within scientific communities.

Current public oceanographic data sets are typically dispersed across numerous data repositories and are stored in a wide array of data formats turning the data integration process into an arduous undertaking. For example, satellite products and numerical model outputs are housed in diverse data repositories and typically delivered in netCDF format [2] following CF-conventions [3]. *In-situ* measurements derived from time-series observations, long-term research expedition programs, or climatologies are generally found at both dedicated websites (e.g., HOT [4], BATS [5], AMT [6], ARGO [7], GO-SHIP [8], World Ocean Atlas [11]) and at different repositories (e.g., BCO-DMO [9], PANGAEA [10]) and are commonly available in a variety of formats, including flat files. Recent efforts link expedition-specific environmental measurements with contemporaneous organism-based molecular data sets from the same expeditions (Planet Microbe [12], TARA [13]). However, systematic expansion of these linkages beyond the expedition data sets remains as a significant technical challenge.

Data integration across data set types is thus becoming increasingly difficult as marine data repositories become stockpiled with varying file formats and naming conventions. Moreover, the absence of a cohesive data distribution strategy that covers the different sectors of oceanography has contributed to the inconsistent data exposure across all research disciplines. Data architectures for more straightforward access to highly heterogeneous data in Earth science is an active research area. Ongoing research efforts in industry include IBM PAIRS [14] and Google Earth Engine [15]. Academic efforts include the PAN-GEO project [16] and efforts within the SciDB community [17]. In this article we present the approach that Simons CMAP is taking to tackle these challenges for the ocean marine-microbes community.

## 2 Identification of community needs

Designing large-scale knowledge infrastructures meant to serve multi-disciplinary, multi-organizational, and geographically distributed scientists has been and will continue to be a critical challenge [18]. The Simons CMAP design principles were guided by a 12-month study of the workflows of a selected number of domain researchers using an approach informed by Participatory Action Research methodology [19, 20]. We employed a mix of ethnographic methods [21], user research methods [22, 23], and Design Thinking [24, 25]. This mixed-method approach provides a non-linear, iterative process to understand community needs, define the parameters of the problem, and identify new solutions through a prototyping and testing process. Ethnographic methods were employed to document and analyze the co-evolution of stakeholder needs and Simons CMAP. Semi-structured interviews were conducted over the course of a year with 51 participants based in 22 academic research laboratories across the United States, Canada, and the United Kingdom. Data captured from meetings and a subset of initial interviews were analyzed and used to design a survey to evaluate how researchers share and preserve their data, which methods and tools are commonly employed, what factors limit use of publicly available data, and specific content or functionalities that would better support research endeavors. A total of 124 participants received and completed the survey.

Survey results revealed obstacles to the synthetic analyses of publicly available data (Fig. 1). Key factors included differences in the temporal and spatial resolution of different data sets; difficulties in finding, retrieving, and subsetting data from different sources; difficulties in integrating data of various formats; and lack of sufficient documentation and metadata associated with the data sets. Desired functionalities included enhanced abilities to share unpublished data among collaborators. The community-engaged research process also identified and helped prioritize which core public data sets should be ingested in the near term into Simons CMAP.

**Figure 1:**
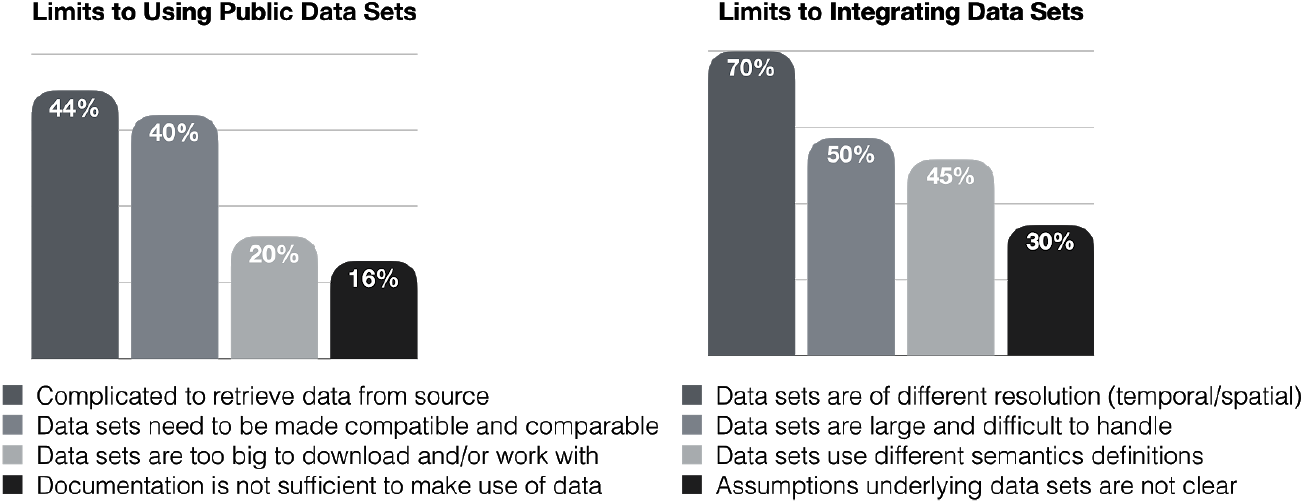
Survey responses: limits to using public data sets and data integration.

## 3 Data harmonization

A key attribute of Simons CMAP is the transformation of publicly available heterogeneous data sets into a common data and metadata structure. To ensure that data sets are Findable, Accessible, Inter-operable, and Reusable (FAIR) [26], the transformed data sets are annotated with a semantic layer that consists of contextual *keywords* about the data set variables. The annotation step is commonly carried out by the data provider or alternatively, by a knowledgeable end-user. The augmented semantic layer makes it possible to identify similar variables registered with different naming conventions. For example, search capabilities are enhanced by the annotation of a variable such as “nitrate” with keywords such as “NO3” or “N+N”, two commonly used alternative designations for nitrate. A web-based validation tool allows individual data providers to confirm their data set adheres to the appropriate formatting requirements and helps to identify potentially erroneous or outlier data points. All annotated data sets are subjected to a final, human-curation step to ensure the proper structure of the data and metadata; a DOI is required for all user-provided data sets as Simons CMAP houses data set copies within its data portal and is not intended as an official master repository. The curated data set copies are ingested into a scalable database system of organized data collections with each data entry indexed by a hashed location and time coordinate. This indexing scheme constitutes the bedrock for efficient data retrieval, visualization, and systematic cross-referencing of the underlying data sets regardless of size or resolution.

The current Simons CMAP database encompasses data sets from a wide array of oceanography sectors selected to help achieve an initial, overarching goal of understanding the structure, function and ecology of marine microbial assemblages. A core component of Simons CMAP is a suite of diverse data sets collected since 2014 on research expeditions conducted through the Simons Collaboration on Ocean Processes and Ecology [27]; a component of the Microbial Oceanography program of the Simons Foundation. These data sets include chemical (e.g., trace elements and organic and inorganic macronutrients), biological (e.g., productivity, particle stochiometry and metabolites, flow cytometry and cell counts, and optical measures), and physical (e.g., temperature, salinity, light) measurements derived from analyzed on-station samples, and from underway and autonomous instruments. The larger oceanographic context is provided by inclusion of global multi-decade remote sensing products (e.g. satellite temperature, chlorophyll, altimetry), several decades of global biogeochemical model estimations (e.g. MIT Darwin [28, 29, 30], Mercator-Pisces [31]), and decades of field measurements (e.g. Argo floats [7], World Ocean Atlas [11], Hawaii Ocean Time series [4]). Numerous additional field expeditions and their associated cruise trajectories conducted over the past decades have been cataloged and ingested in the database, a process that will be actively continued. Figure 2 illustrates a selected list of public data sets that are harmonized and ingested in the Simons CMAP database. The most updated list of datasets can be found at the Simons CMAP website [32].

**Figure 2:**
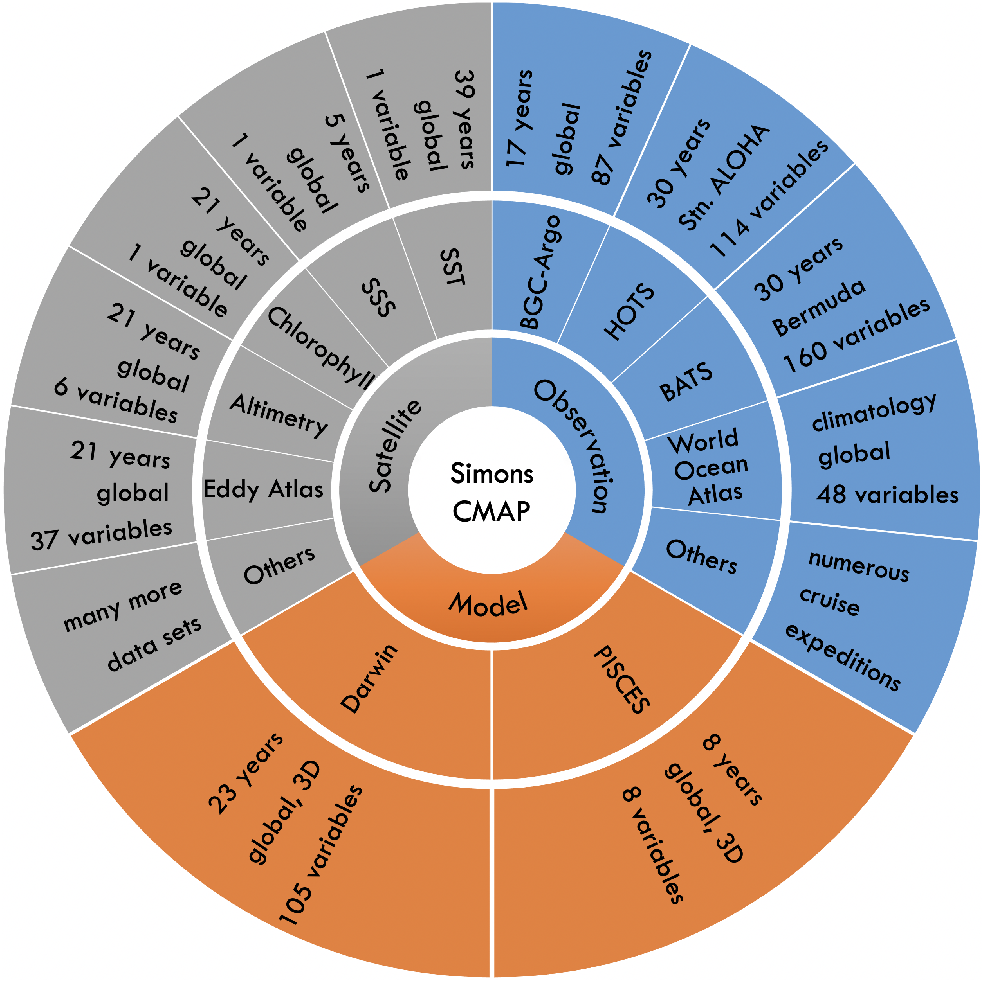
A schematic illustration showing a selected list of cross-discipline ocean data sets hosted by Simons CMAP. Data sets representing direct field observations are in blue, remote sensing data sets are in gray, and model estimates are organized in the orange section.

The Simons CMAP database accepts public and open-source data sets with explicit time and location annotations. We invite the Oceanography research community to submit their data sets via our data submission portal [33] where the data harmonization and ingestion process is initiated through a series of step-by-step operations. Once ingested, the data set can be visualized using the plotting services and can be integrated with any other archived data set.

## 4 System overview

Over the past decade, Earth science data management systems have progressively adopted advanced technologies underpinning the Big Data ecosystems. While these new advancements open the door to the efficient handling of massive data collections, they also raise the technical skills required to access and analyze the amassed data. Simons CMAP is designed to lower the required technical competencies by abstracting away the details introduced by the underlying technologies. Figure 3 depicts a simplified schematic of the Simons CMAP process-flow illustrating how the complexities associated with data preparation and management technologies are decoupled from the application layer where the end-user interacts with data.

**Figure 3:**
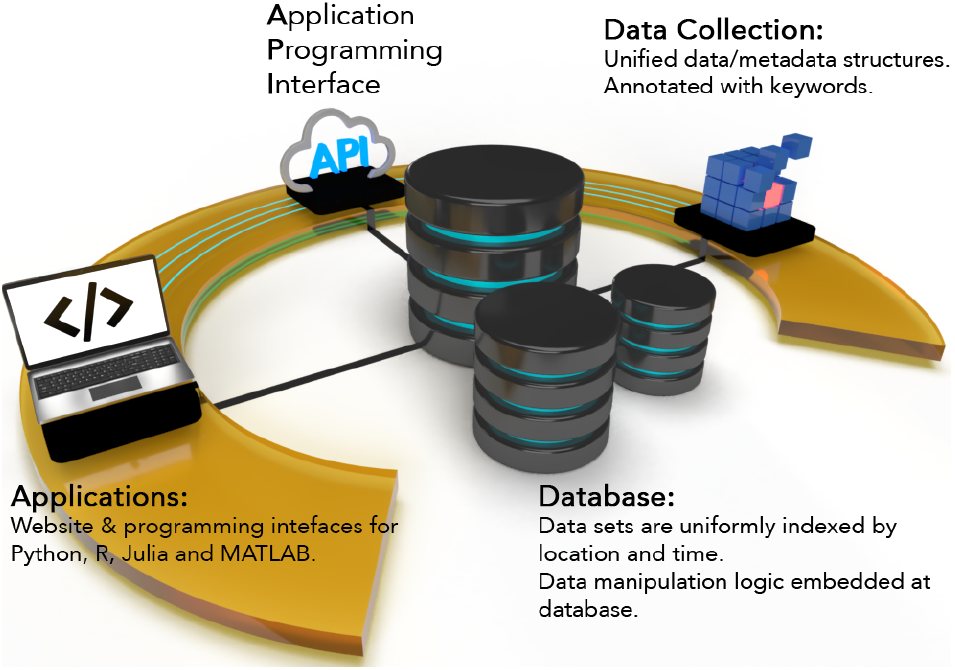
A schematic overview of the Simons CMAP architecture. Data sets with miscellaneous formats are collected and transformed into a common structure. The transformed data sets along with their associated metadata are loaded into database systems where data is indexed by location and time. A cloud-based API exposes the database contents to the end-user and external applications abstracting away the details associated with the underlying technologies. The applications provide graphical and programming interfaces to retrieve, visualize, and analyze the data.

The data pipeline begins with the “collection” step where the collected data sets are curated and harmonized by location and time prior to being transferred to the database systems [34]. Each data set is populated in a *table* where columns represent the data set *variables*. The database acts as the central component as it hosts both the data and the data extraction logic. Integrating the extraction logic into the heart of the database enables two key features: **a)** the technical process of data retrieval or co-localization of two different data sets is substantially simplified as it is handled internally and **b)** Simons CMAP becomes a language-agnostic product allowing researchers to retrieve the data with their [programming] language of choice by invoking these predefined database-level services. A cloud-based REST Application Programming Interface (API) [35, 36] operates as a gateway to the database resources and handles data requests issued by the downstream applications. The outermost component of the architecture is comprised of an application suite developed to expose the Simons CMAP services to the end-users. To date, these applications include an interactive web interface that allows for data submission, access, and visualization [32] plus a series of programming interfaces supporting Python [37], R [38], Matlab [39] and Julia [40] languages. These documented programming interfaces [41] provide easy access to the Simons CMAP services and make it possible to conduct controlled and reproducible research studies. Listing 1 shows a simple script that uses the Simons CMAP Python package (`pycmap`) to retrieve a subset of salinity measurements (`variable=’argo_merge_salinity_adj’`) stored at the Argo data set (`table=’tblArgoMerge_REP’`). Data is returned in form of a DataFrame object; a widely-accepted data structure within the scientific computing community with extensive built-in analytical features [42]. In addition to extracting subsets of data, the programming interfaces present methods for data visualization, analytics such as aggregation along time and space axes (e.g. time-series, depth profiles), computing dataset-specific climatology with custom time-frames (e.g. weekly, monthly, quarterly climatologies), and co-localization services (see documentation for more details and examples [43]).

### Listing 1

Example Python code showing how to retrieve a data set slice. `space_time` is one of the Simons CMAP API methods that returns subset of a variable delimited by spatio-temporal constrains. A variable is identified by its name (`variable=’argo_merge_salinity_adj’`) and the name of the table where it resides (`table=’tblArgoMerge_REP’`). Refer to the online documentation [43] for a complete list of API methods and sample codes. Note that using Simons CMAP programming interfaces requires an API Key which can be obtained freely from the website.

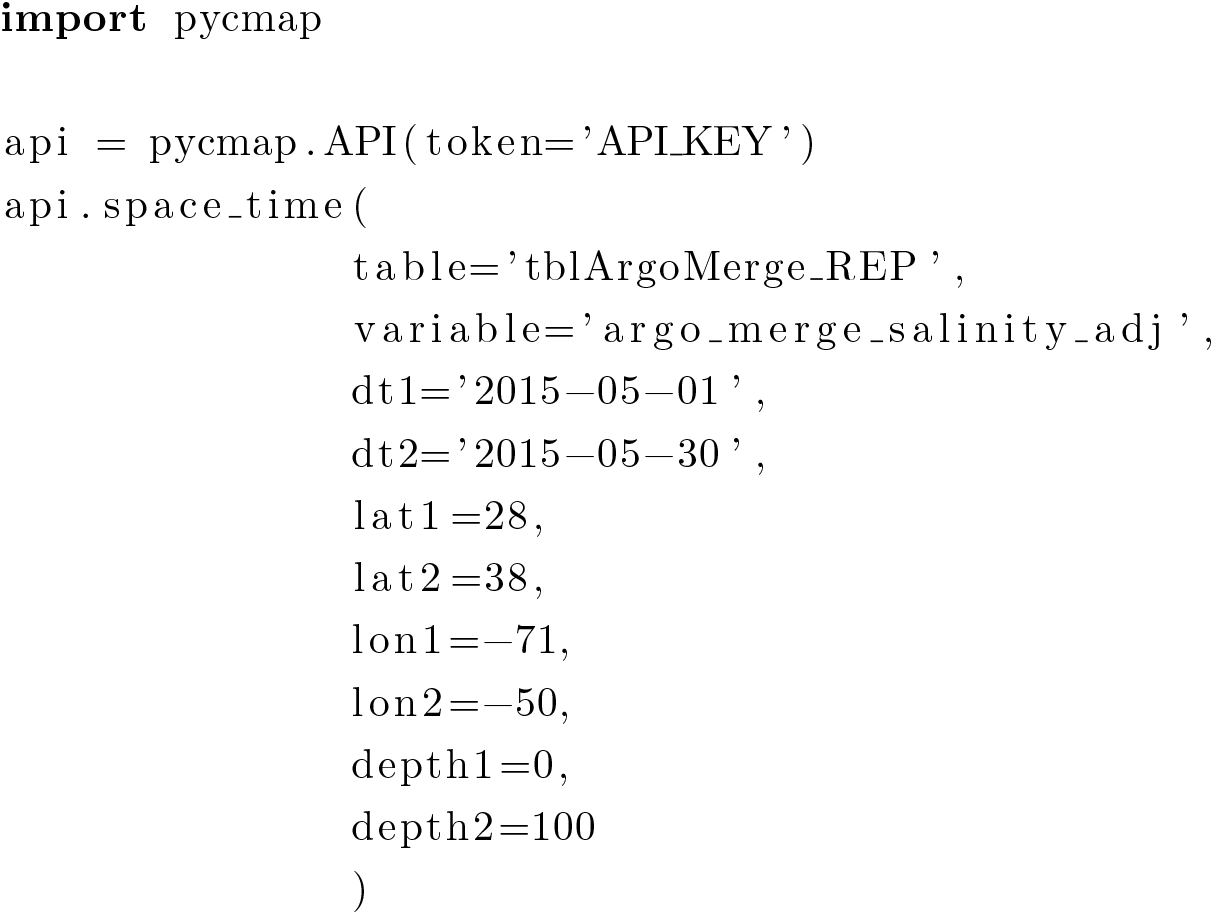

To extract a variable slice using the programming interfaces, one needs to know the exact variable name and the name of the table where it is hosted. The latest list of variable and table names is accessible at the website’s catalog page (https://simonscmap.com/catalog) which offers an interactive alternative for data retrieval. Moreover, the website provides the user with an intuitive plotting toolset for creating a range of data visualizations without prior knowledge of the exact variable and table names. Including multiple interfaces is a feature incorporated in response to user interviews, which identified a wide range in technical expertise among users. The combination of a public web interface and a documented public API is designed to map to these different needs identified in community discussions.

## 5 Data Integration Example

Marine observations reflect an intrinsically complex system that is under the influence of interwoven physical, chemical, and biological dynamics. Arguably, one of the most effective approaches to understanding these processes is conducting synthetic analyses where the field observations are joined with independent ancillary data sets such as environmental parameters to leverage their collective insight. However, collecting and integrating multiple different massive marine data sets can be a substantially challenging and time-consuming process as they are obtained from various sources with inconsistent data structures. Here we describe a multi-variate analysis example where we compile observations of specific phytoplankton groups archived within Simons CMAP and co-localize them with the adjacent environmental conditions derived from independent data sets. Since all Simons CMAP data sets are uniformly harmonized, the entire process of collecting and integrating multiple massive global data sets with different dimensions and resolutions can be summarized into short code blocks developed using the Simons CMAP programming interfaces.

The planet’s oceans are teeming with diverse microbial communities that drive the biogeochemical processes delineating the global biogeochemical cycles [44]. Marine phytoplankton account for approximately 50% of global net primary production and therefore have a critical impact on the production and cycling of nutrients and organic matters [45]. The most abundant members of the phytoplankton are the picophytoplankton with cell diameters less than ≈ 3 *μ*m. They are distributed throughout lighted waters of the global oceans. The picophytoplankton are composed of *Prochlorococcus* and *Synechococcus*, two important cyanobacteria, and the pico-eukaryotes, a mixture of different eukaryotic pico-phytoplankton groups. *Prochlorococcus* is the most abundant photosynthetic organism on the planet and is the numerically dominant phototroph in tropical and subtropical oceans, with key contributions to the ecosystems and global biochemical cycles [46].

The Simons CMAP database hosts more than three decades of global measurements of picophytoplankton characteristics in a unified format. These data sets are derived from multi-decade research programs such as the Atlantic Meridional Transect (AMT) [6], global compilations of cruise expeditions and time-series programs [47, 48] and high-resolution observations recorded using SeaFlow, a continuous shipboard flow cytometer [49]. The compiled data set for this case study consists of more than 163,000 flow cytometry-based measurements of volumetric cell abundances of *Prochlorococcus*, *Synechococcus*, and pico-eukaryotes from the top 5 meters of the global oceans during an approximately 35-year period (Fig. 4). The complete list of gathered data sets can be found in the supplementary materials (Table S1).

**Figure 4:**
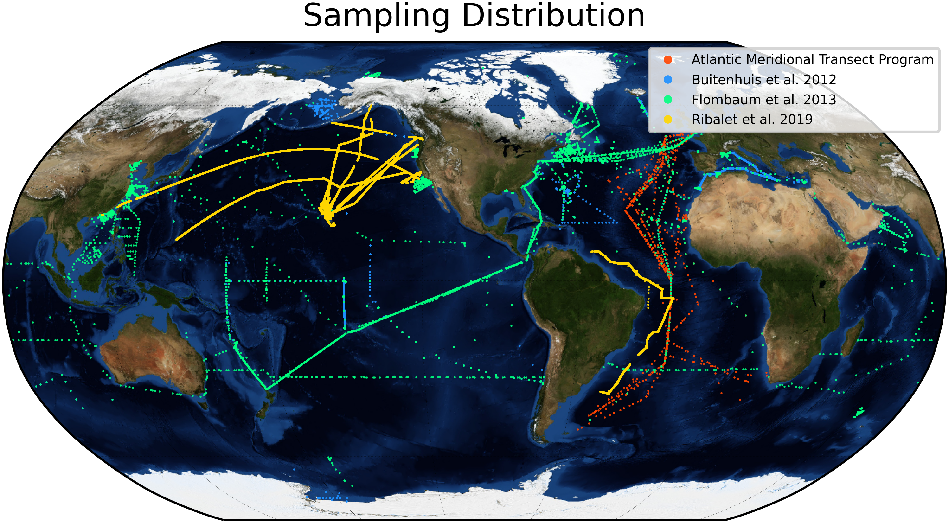
Geographic distribution of collected Cyanobacteria observations. The collected data sets are results of more than 35 years of field expeditions involving 16 cruise expeditions conducted by the Atlantic Meridional Transect Program [6], 50 cruise expeditions by the Sea Flow Program [49], and decades worth of time-series and cruise expeditions compiled by [47, 48]. The samples are restricted to the surface waters (≈ 5 m depth).

The Simons CMAP database is structured to handle the process of co-localizing *in-situ* measurements with environmental variables from diverse data sets, based on user-defined temporal and spatial tolerance parameters (Fig. 5). Here, each flow cytometry-based abundance measurement for the pico-phytoplankton (*Prochlorococcus*, *Synechococcus*, pico-eukaryotes) was co-localized with a selected list of 20 contemporaneous environmental parameters including temperature, salinity, nutrient concentrations, altimetry, and surface geostrophic currents derived from multi-decade global remote-sensing and numerical model data sets (see supplementary materials Table S2). The tolerance parameters used for the co-localization procedure varied depending on the resolution of the underlying environmental data set, but typically they were set to ±0.25*°* for the meridional and zonal directions, ±5 m for the vertical direction, and the temporal tolerance was generally set to ±1 day or to ±4 days for weekly averaged data sets. A monthly climatology value was used when observation data points fell outside the temporal coverage of a given environmental data set. The outcome of this procedure is a large compilation of archived sea surface Cyanobacteria cell abundance observations that are integrated with contemporaneous environmental variabales. This data set can then be used to quantify the correlation between the observations and environmental factors, and consequently to conduct more detailed statistical analyses. As a simple example, Fig. 6 illustrates the large-scale correlation between *Prochlorococcus* cell abundances and environmental variables such as temperature and dissolved phosphate concentrations (panels **a** and **b**), highlighting Simons CMAP capabilities in integrating data sets from various sectors of Oceanography (panel **c**).

**Figure 5:**
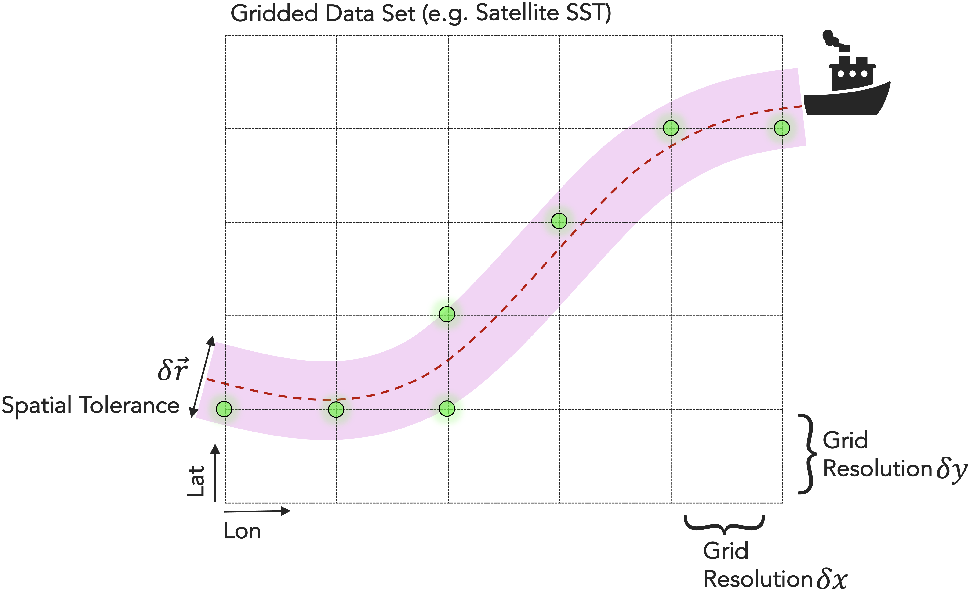
A schematic illustration demonstrating how a cruise trajectory is co-localized with a gridded product such as satellite Sea Surface Temperature (SST) using user-defined spatio-temporal tolerance parameters. Satellite temperature measurements are linked with the cruise data set if they fall within the spatial tolerance parameter 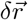 (zonal and meridional tolerances). If more than one temperature data point match the conditions, their mean value is linked with the cruise data set. The green dots represent the matched grid data points with the cruise trajectory according to the tolerance parameters. Similar tolerance parameters can be defined along the depth and time axes (not shown in the figure).

**Figure 6:**
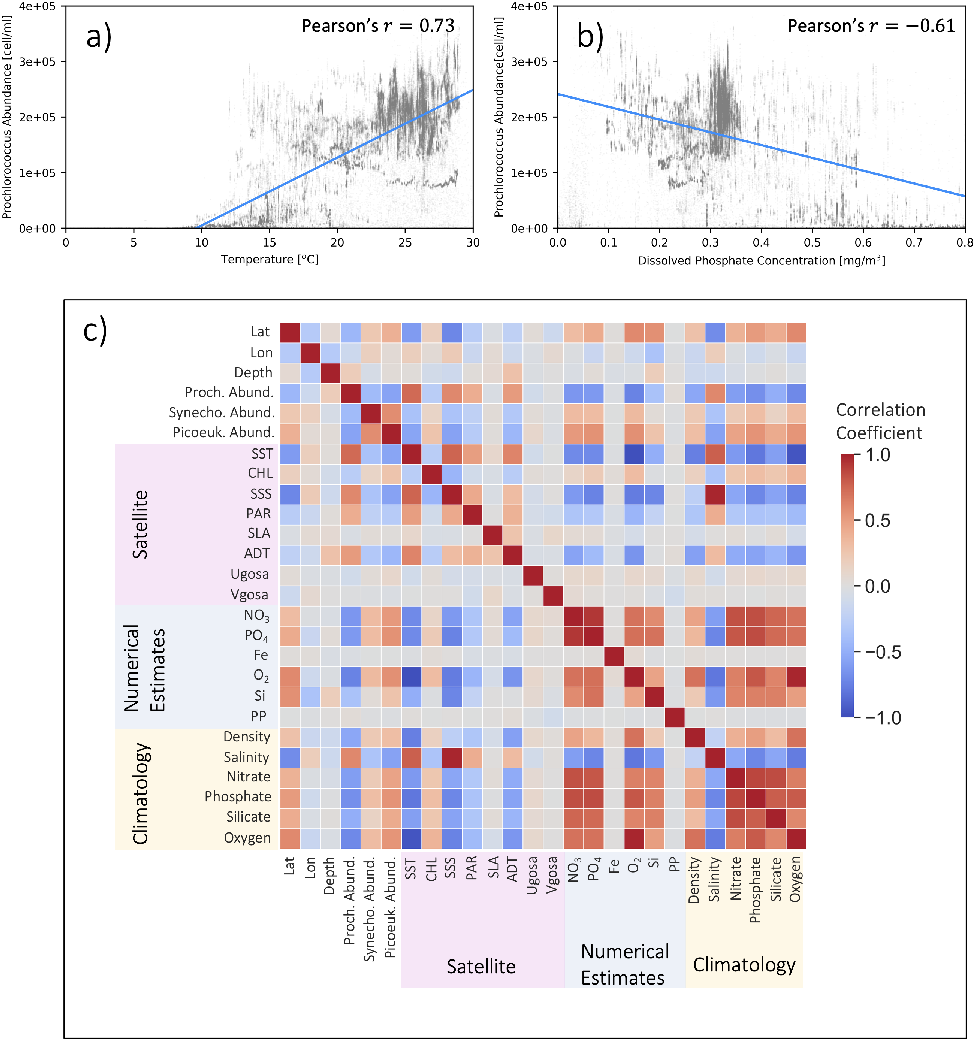
Panels **a** and **b** show the compiled measurements of sea surface *Prochlorococcus* cell abundances as a function of remotely-sensed temperature and phosphate concentrations estimated by a numerical model [31]. Panel **c** presents pairwise Pearson’s correlation coefficients computed between the integrated environmental variables and the Cyanobacteria cell abundance observations. For more details about the environmental data sets refer to supplementary materials (Table S2). Uniform data harmonization schemes applied across the Simons CMAP database make it possible to interconnect *in-situ* observations with satellite data and model outputs.

In the absence of Simons CMAP, one would instead need to first identify and collect decades of global observations of pico-phytoplankton abundances, a substantially time-consuming process as these data sets are scattered across the web and research literature. The collected data sets would then need to be co-localized with ancillary data sets (e.g. decades of global satellite data) with varying resolutions and file formats, a laborious and challenging undertaking. In contrast, due to the local storage of these data sets in the Simons CMAP database and harmonization of the data by location and time, the entire task of collecting and integrating the organism abundances with the environmental factors can be accomplished using short and simple scripts saving a substantial amount of data preparation time. The scripts to collect and integrate data sets and the final compiled data set are publicly accessible [50].

## 6 Conclusion

Simons CMAP is designed to alleviate current barriers to utilizing oceanographic data sets that can be difficult to find, may be massive in size, and are typically released in a variety of file formats and resolutions. Our strategy is to collect, curate, and construct a database that hosts a wide spectrum of data sets from all sectors of Oceanography in a unifying data structure, combined with simple interfaces to retrieve and analyze the data. The uniform harmonization of data entries by location and time substantially facilitates the process of integrating data sets which in turn can lead to discovery of intriguing scientific patterns. Many ideas in environmental research involve characterizing the spatiotemporal relations amongst diverse properties. Simons CMAP creates a unified system that accelerates the ability to explore ideas and hypotheses against highly heterogeneous and diverse mixes of data. Key to this is a unified database architecture and a focus on an interface that aligns with the target research community.

An essential component of the genesis of Simons CMAP was surveys and interviews of diverse researchers to gain insights into how, and under what circumstances, domain scientists across various disciplines of Oceanography discover, acquire, share, and use data as part of their collaborative research work. Such iterative and ongoing involvement of the larger community is essential to creation of data research infrastructures appropriate to support cutting edge research. Development of Simons CMAP will continue to be informed by ongoing usability tests and user interviews.

The Simons CMAP project is envisioned to progressively increase in size and diversity into the future, with a focus on organizing the currently widely dispersed data from research cruises and placing these data within the larger context of satellite products and output from numerical models. The longer-term goal is to develop the infrastructure necessary to collaborate with other marine data initiatives to construct a unified network of interconnected and distributed databases. Development of standard APIs that interconnect these databases with compilations of environmental data such as those found in Simons CMAP will significantly contribute to the progress of multidisciplinary studies.

## Data availability

More than three decades of Cyanobacteria cell abundance observations are compiled and co-localized with a variety of environmental variables. The code developed to collect and integrate, as well as the final generated data set is publicly available [50].

## Supplementary data

Supplementary material is available at the online version of the manuscript.

## Acknowledgements

We thank all research laboratories supported by Simons Foundation Life Science program that have provided their data sets to Simons CMAP database. We thank Aditya Mishra, Sangwon Hyun, Christian Müller, and Jacobe Bien for developing the R programming interface. We thank Megan Schatz for proof-reading the system’s documentations and helping N.R.H to organize and process a subset of data sets, and Hunter Hadaway for Simons CMAP web design. We also thank the following organizations for providing public marine data sets: **Marine Copernicus:** “*Data was provided by: E.U. Copernicus Marine Service Information*”, **Argo Floats Program:** “*These data were collected and made freely available by the International Argo Program and the national programs that contribute to it. (http://www.argo.ucsd.edu , http://argo.jcommops.org). The Argo Program is part of the Global Ocean Observing System*”. Likewise we thank **AVISO program** (https://www.aviso.altimetry.fr/), **World Ocean Atlas** (https://www.nodc.noaa.gov/OC5/woa18/), and **Jet Propulsion Laboratory** (https://podaac-opendap.jpl.nasa.gov/). This work was supported by a grant from the Simons Foundation (Award ID 549945 to E.V.A).

## Author contributions

M.D.A led the project, architected the database systems, developed Python, MATLAB, and Julia programming interfaces, wrote API documentations, designed and build system’s hardware infrastructure, and wrote this manuscript with contributions from all authors. N.R.H and R.L.M contributed to data collection, curation, ingestion, and published documentations. M.D contributed by implementing the website. T.C.B. contributed by acting as the project coordinator. A.N and C.P.L conducted surveys to identify system core functionalities and consequently performed usability tests. C.N.H contributed to the early formation of the project architecture. As the project director, E.V.A supervised the program.

## References

[1] Pendleton L. H. , Beyer H., Estradivari, Grose S. O., Hoegh-Guldberg O., Karcher D. B., Kennedy E., Llewellyn L., Nys C., Shapiro A., Jain R., Kuc K., Leatherland T., O’Hainnin K., Olmedo G., Seow L., and Tarsel M. 2019. Disrupting data sharing for a healthier ocean. ICES Journal of Marine Science, 76(6): 1415–1423, https://doi.org/10.1093/icesjms/fsz068.

[2] Unidata: Network Common Data Form (NetCDF). Boulder, CO: UCAR/Unidata Program Center, https://doi.org/10.5065/D6H70CW6.

[3] Eaton B. , Gregory J., Drach B., Taylor K., Hankin S., Blower J., Caron J., Signell R., Bentley P., Rappa G., Höck H., Pamment A., Juckes M., and Raspaud M. NetCDF Climate and Forecast (CF) Metadata Conventions. http://cfconventions.org/.

[4] Web Access: http://hahana.soest.hawaii.edu/hot/.

[5] Web Access: http://bats.bios.edu/.

[6] Robins D. B. , and Aiken J. 1996. The Atlantic Meridional Transect: an oceanographic research programme to investigate physical, chemical, biological and optical variables of the Atlantic Ocean. Underwater Technology, 21(4): 8–14, https://doi.org/10.3723/175605496783328529.

[7] Argo float data and metadata from the Global Data Assembly Center (Argo GDAC), SEANOE: 2019. https://doi.org/10.17882/42182.

[8] Web Access: https://www.go-ship.org/.

[9] Web Access: https://www.bco-dmo.org/.

[10] Web Access: https://www.pangaea.de/.

[11] Boyer T. P. , Antonov J. I., Baranova O. K., Coleman C., Garcia H. E., Grodsky A., Johnson D. R., Locarnini R. A., Mishonov A. V., O’Brien T. D., Paver C. R., Reagan J. R., Seidov D., Smolyar I. V., and Zweng M. M. 2013. World Ocean Database 2013, NOAA Printing Office, Silver Spring, MD, https://data.nodc.noaa.gov/woa/WOD13/DOC/wod13_intro.pdf.

[12] Ponsero A. J. , Bomhoff M., Blumberg K., Youens-Clark K., Herz N. M., Wood-Charlson E. M., Delong E. F., and Hurwitz B. L. 2020. Planet Microbe: a platform for marine microbiology to discover and analyze interconnected omics and environmental data. Nucleic Acids Research, 49: D792–D802 https://doi.org/10.1093/nar/gkaa637.

[13] Web Access: https://oceans.taraexpeditions.org/en/.

[14] Lu S. , Shao X., Freitag M., Klein L. J., Renwick J., Marianno F. J., Albrecht C., Hamann H. F. 2016. IBM PAIRS curated big data service for accelerated geospatial data analytics and discovery. 2016 IEEE International Conference on Big Data (Big Data), pp. 2672–2675, https://doi.org/10.1109/BigData.2016.7840910.

[15] Gorelick N. , Hancher M., Dixon M., Ilyushchenko S., Thau D., Moore R. 2017. Google Earth Engine: Planetary-scale geospatial analysis for everyone. Remote sensing of Environment, 202: 18–27, https://doi.org/10.1016/j.rse.2017.06.031.

[16] Eynard-Bontemps G. , Abernathey R., Hamman J., Ponte A., Rath W. 2019. The PANGEO Big Data Ecosystem and its use at CNES, pp. 49–52, https://doi.org/10.2760/848593.

[17] Appel M. , Lahn F., Buytaert W., Pebesma E. 2018. Open and scalable analytics of large Earth observation datasets: From scenes to multidimensional arrays using SciDB and GDAL. ISPRS journal of photogrammetry and remote sensing, 138: 47–56, https://doi.org/10.1016/j.isprsjprs.2018.01.014.

[18] Atkins, D. E. , Droegemeier, K. K., Feldman, S. I., et al. 2003. Revolutionizing science and engineering through cyberinfrastructure: Report of the National Science Foundation blue-ribbon advisory panel on cyberinfrastructure. National Science Foundation, Washington, DC, https://www.nsf.gov/cise/sci/reports/atkins.pdf.

[19] McTaggart, R. 1997. Participatory action research: International contexts and consequences, State University of New York Press, Albany, NY.

[20] Kemmis, S., and McTaggart, R. 2005. Participatory Action Research: Communicative Action and the Public Sphere. In The Sage handbook of qualitative research, 3rd ed., pp. 559–603. Sage Publications Ltd, Thousand Oaks, CA.

[21] Marcus, G. E. 1995. Ethnography in/of the world system: The emergence of multi-sited ethnography. Annual review of anthropology, 24: 95–117, https://doi.org/10.1146/annurev.an.24.100195.000523.

[22] Baxter, K., Courage, C., and Caine, K. 2015. Understanding Your Users: A Practical Guide to User Research Methods, Morgan Kaufmann Publishers Inc, https://doi.org/10.1016/C2013-0-13611-2.

[23] Rohrer, C. 2014. When to use which user-experience research methods. Nielsen Norman Group: 1–7, https://www.nngroup.com/articles/which-ux-research-methods/.

[24] Dorst, K. 2011. The core of ‘design thinking’ and its application. Design studies, 32: 521–532, https://doi.org/10.1016/j.destud.2011.07.006.

[25] Plattner, H., Meinel, C., and Leifer, L. 2012. Design Thinking Research: Studying Co-creation in Practice, Springer, https://doi.org/10.1007/978-3-642-21643-5.

[26] Wilkinson M. D. , Dumontier M, Aalbersberg I. J. J., et al. 2016. The FAIR Guiding Principles for scientific data management and stewardship. Scientific Data, 3: 160018, https://doi.org/10.1038/sdata.2016.18.

[27] Web Access: https://www.simonsfoundation.org/life-sciences/microbial-oceanography/simons-collaboration-on-ocean-processes-and-ecology/.

[28] Dutkiewicz S. , Hickman A. E., Jahn O., Gregg W. W., Mouw C. B., and Follows M. J. 2015. Capturing optically important constituents and properties in a marine biogeochemical and ecosystem model. Biogeoscience, 12: 4447–4481, https://doi.org/10.5194/bg-12-4447-2015.

[29] Ward B. A. , Dutkiewicz S., Jahn O., and Follows M. J. 2012. A size-structured food-web model for the global ocean. Limnology and Oceanography, 57: 1877–1891, https://aslopubs.onlinelibrary.wiley.com/doi/abs/10.4319/lo.2012.57.6.1877.

[30] Web Access: http://darwinproject.mit.edu/.

[31] Aumont O. , Ethé C., Tagliabue A., Bopp L., and Gehlen M. 2015. PISCES-v2: an ocean biogeochemical model for carbon and ecosystem studies. Geoscitific Model Development, 8: 2465–2513, https://doi.org/10.5194/gmd-8-2465-2015.

[32] Web Access: https://simonscmap.com.

[33] Web Access: https://simonscmap.com/datasubmission.

[34] Web Access: https://github.com/simonscmap/DB

[35] Fielding R. T. 2000. Architectural Styles and the Design of Network based Software Architecture. Ph.D. Thesis. University of California, Irvine. 76–85, https://www.ics.uci.edu/~fielding/pubs/dissertation/fielding_dissertation.pdf.

[36] Web Access: https://github.com/simonscmap/cmap-node-api.

[37] Web Access: https://github.com/simonscmap/pycmap, https://doi.org/10.5281/zenodo.3387777.

[38] Web Access: https://github.com/simonscmap/cmap4r.

[39] Web Access: https://github.com/simonscmap/matcmap.

[40] Web Access: https://github.com/simonscmap/CMAP.jl.

[41] Web Access: https://cmap.readthedocs.io.

[42] McKinney W. 2010. Data Structures for Statistical Computing in Python. Proceedings of the 9th Python in Science Conference, 51–56, https://doi.org/10.25080/Majora-92bf1922-00a.

[43] Web Access: https://cmap.readthedocs.io/en/latest/user_guide/API_ref/api_ref.html

[44] Falkowski P. G. , Fenchel T., Delong E. F. 2008. The microbial engines that drive Earth’s biogeochemical cycles. Science, 320: 1034–1039, https://doi.org/10.1126/science.1153213.

[45] Field C. B. , Behrenfeld M. J., Randerson J. T., Falkowski P. 1998. Primary production of the biosphere: Integrating terrestrial and oceanic components. Science, 281: 237–240, https://doi.org/10.1126/science.281.5374.237.

[46] Moore L. R. , Chisholm S. W. 1999. Photophysiology of the marine cyanobacterium Prochlorococcus: Ecotypic differences among cultured isolates. Limnology and Oceanography, 44: 628–638, https://doi.org/10.4319/lo.1999.44.3.0628.

[47] Flombaum P. , Gallegos J. L., Gordillo R. A., Rincón J., Zabala L. L., Jiao N., Karl D. M., Li W. K. W., Lomas M. W., Veneziano D., Vera C. S., Vrugt J. A., Martiny A. C. 2013. Present and future global distributions of the marine Cyanobacteria Prochlorococcus and Synechococcus. PNAS, 24: 9824–9829, https://doi.org/10.1073/pnas.1307701110.

[48] Buitenhuis E. T. , Li K. W. W., Vaulot D., Lomas M. W., Landry M. R., Partensky F., Karl D. M., Ulloa O., Campbell L., Jacquet S., Lantoine F., Chavez F. P., Macias D., Gosselin M., McManus G. B. 2012. Global distributions of picophytoplankton abundance and biomass - Gridded data product (NetCDF) - Contribution to the MAREDAT World Ocean Atlas of Plankton Functional Types, Earth System Science Data, 4, 37–46, https://doi.org/10.5194/essd-4-37-2012.

[49] Ribalet F. , Bertiaume C., Hynes A., Swalwell J., Carlson M., Clayton S., Hennon G., Poirier Ca., Shimabukuro E., White A., Armbrust E. V. SeaFlow data 1.0: 2019. high-resolution abundance, size and biomass of small phytoplankton in the North Pacific. Scientific Data, 6: 277, https://doi.org/10.6084/m9.figshare.10110755.

[50] Askezari M. D. , Hagen N. R., Denholtz M., Neang A., P. Lee C., Armbrust E. V. 2020. Compiled Data Set of Sea Surface Prochlorococcus, Synechococcus, and pico-eukaryotes Using Simons CMAP Database, https://doi.org/10.5281/zenodo.4108150.

