## Supplementary Materials for "Simons Collaborative Marine Atlas Project (Simons CMAP): an open-source portal to share, visualize and analyze ocean data"

Table S1: List of cyanobacteria data sets collected for the case study (Sec. 5). The "Table" column refers to the database table name where the data set is stored and the "Variable" column specifies the exact variable name in the Simons CMAP database.

| | Table | Variable | Dataset Page |
|---|---|---|---|
| 1 | tblSeaFlow | abundance_prochloro | https://simonscmap.com/catalog/datasets/all_SeaFlow_cruises |
| 2 | tblSeaFlow | abundance_synecho | https://simonscmap.com/catalog/datasets/all_SeaFlow_cruises |
| 3 | tblSeaFlow | abundance_picoeuk | https://simonscmap.com/catalog/datasets/all_SeaFlow_cruises |
| 4 | tblFlombaum | prochlorococcus_abundance_flombaum | https://simonscmap.com/catalog/datasets/Flombaum |
| 5 | tblFlombaum | synechococcus_abundance_flombaum | https://simonscmap.com/catalog/datasets/Flombaum |
| 6 | tblGlobal_PicoPhytoPlankton | prochlorococcus_abundance | https://simonscmap.com/catalog/datasets/Global_Picophytoplankton |
| 7 | tblGlobal_PicoPhytoPlankton | synechococcus_abundance | https://simonscmap.com/catalog/datasets/Global_Picophytoplankton |
| 8 | tblGlobal_PicoPhytoPlankton | picoeukaryote_abundance | https://simonscmap.com/catalog/datasets/Global_Picophytoplankton |
| 9 | tblJR19980514_AMT06_Flow_Cytometry | prochlorococcus_abundance_P701A90Z_Zubkov | https://simonscmap.com/catalog/datasets/JR19980514_AMT06_flow_cytometry |
| 10 | tblJR19980514_AMT06_Flow_Cytometry | synechococcus_abundance_P700A90Z_Zubkov | https://simonscmap.com/catalog/datasets/JR19980514_AMT06_flow_cytometry |
| 11 | tblJR19980514_AMT06_Flow_Cytometry | picoeukaryotic_abundance_PYEUA00A_Zubkov | https://simonscmap.com/catalog/datasets/JR19980514_AMT06_flow_cytometry |
| 12 | tblJR20030512_AMT12_Flow_Cytometry | prochlorococcus_abundance_P701A90Z_Zubkov | https://simonscmap.com/catalog/datasets/JR20030512_AMT12_flow_cytometry |
| 13 | tblJR20030512_AMT12_Flow_Cytometry | synechococcus_abundance_P700A90Z_Zubkov | https://simonscmap.com/catalog/datasets/JR20030512_AMT12_flow_cytometry |
| 14 | tblJR20030512_AMT12_Flow_Cytometry | picoeukaryotic_abundance_PYEUA00A_Zubkov | https://simonscmap.com/catalog/datasets/JR20030512_AMT12_flow_cytometry |
| 15 | tblJR20030910_AMT13_Flow_Cytometry | prochlorococcus_abundance_P701A90Z_Zubkov | https://simonscmap.com/catalog/datasets/JR20030910_AMT13_flow_cytometry |
| 16 | tblJR20030910_AMT13_Flow_Cytometry | synechococcus_abundance_P700A90Z_Zubkov | https://simonscmap.com/catalog/datasets/JR20030910_AMT13_flow_cytometry |
| 17 | tblJR20030910_AMT13_Flow_Cytometry | picoeukaryotic_abundance_PYEUA00A_Zubkov | https://simonscmap.com/catalog/datasets/JR20030910_AMT13_flow_cytometry |
| 18 | tblJR20040428_AMT14_Flow_Cytometry | prochlorococcus_abundance_P701A90Z_Zubkov | https://simonscmap.com/catalog/datasets/JR20040428_AMT14_flow_cytometry |
| 19 | tblJR20040428_AMT14_Flow_Cytometry | synechococcus_abundance_P700A90Z_Zubkov | https://simonscmap.com/catalog/datasets/JR20040428_AMT14_flow_cytometry |
| 20 | tblJR20040428_AMT14_Flow_Cytometry | picoeukaryotic_abundance_PYEUA00A_Zubkov | https://simonscmap.com/catalog/datasets/JR20040428_AMT14_flow_cytometry |
| 21 | tblD284_AMT15_Flow_Cytometry | prochlorococcus_abundance_P701A90Z_Zubkov | https://simonscmap.com/catalog/datasets/D284_AMT15_flow_cytometry |
| 22 | tblD284_AMT15_Flow_Cytometry | synechococcus_abundance_P700A90Z_Zubkov | https://simonscmap.com/catalog/datasets/D284_AMT15_flow_cytometry |
| 23 | tblD294_AMT16_Flow_Cytometry | prochlorococcus_abundance_P701A90Z_Tarran | https://simonscmap.com/catalog/datasets/D294_AMT16_flow_cytometry |
| 24 | tblD294_AMT16_Flow_Cytometry | synechococcus_abundance_P700A90Z_Tarran | https://simonscmap.com/catalog/datasets/D294_AMT16_flow_cytometry |
| 25 | tblD294_AMT16_Flow_Cytometry | picoeukaryotic_abundance_PYEUA00A_Tarran | https://simonscmap.com/catalog/datasets/D294_AMT16_flow_cytometry |
| 26 | tblD299_AMT17_Flow_Cytometry | prochlorococcus_abundance_P701A90Z_Zubkov | https://simonscmap.com/catalog/datasets/D299_AMT17_flow_cytometry |
| 27 | tblD299_AMT17_Flow_Cytometry | synechococcus_abundance_P700A90Z_Zubkov | https://simonscmap.com/catalog/datasets/D299_AMT17_flow_cytometry |
| 28 | tblD299_AMT17_Flow_Cytometry | picoeukaryotic_abundance_PYEUA00A_Zubkov | https://simonscmap.com/catalog/datasets/D299_AMT17_flow_cytometry |
| 29 | tblJR20081003_AMT18_flow_cytometry | prochlorococcus_abundance_P701A90Z_Tarran | https://simonscmap.com/catalog/datasets/JR20081003_AMT18_flow_cytometry |
| 30 | tblJR20081003_AMT18_flow_cytometry | synechococcus_abundance_P700A90Z_Tarran | https://simonscmap.com/catalog/datasets/JR20081003_AMT18_flow_cytometry |
| 31 | tblJR20081003_AMT18_flow_cytometry | picoeukaryotic_abundance_PYEUA00A_Tarran | https://simonscmap.com/catalog/datasets/JR20081003_AMT18_flow_cytometry |
| 32 | tblJC039_AMT19_flow_cytometry | prochlorococcus_abundance_P701A90Z_Tarran | https://simonscmap.com/catalog/datasets/JC039_AMT19_flow_cytometry |
| 33 | tblJC039_AMT19_flow_cytometry | synechococcus_abundance_P700A90Z_Tarran | https://simonscmap.com/catalog/datasets/JC039_AMT19_flow_cytometry |
| 34 | tblJC039_AMT19_flow_cytometry | picoeukaryotic_abundance_PYEUA00A_Tarran | https://simonscmap.com/catalog/datasets/JC039_AMT19_flow_cytometry |
| 35 | tblJC053_AMT20_flow_cytometry | prochlorococcus_abundance_P701A90Z_Tarran | https://simonscmap.com/catalog/datasets/JC053_AMT20_flow_cytometry |
| 36 | tblJC053_AMT20_flow_cytometry | synechococcus_abundance_P700A90Z_Tarran | https://simonscmap.com/catalog/datasets/JC053_AMT20_flow_cytometry |
| 37 | tblJC053_AMT20_flow_cytometry | picoeukaryotic_abundance_PYEUA00A_Tarran | https://simonscmap.com/catalog/datasets/JC053_AMT20_flow_cytometry |
| 38 | tblD371_AMT21_flow_cytometry | prochlorococcus_abundance_P701A90Z_Tarran | https://simonscmap.com/catalog/datasets/D371_AMT21_flow_cytometry |
| 39 | tblD371_AMT21_flow_cytometry | synechococcus_abundance_P700A90Z_Tarran | https://simonscmap.com/catalog/datasets/D371_AMT21_flow_cytometry |
| 40 | tblD371_AMT21_flow_cytometry | picoeukaryotic_abundance_PYEUA00A_Tarran | https://simonscmap.com/catalog/datasets/D371_AMT21_flow_cytometry |
| 41 | tblJC079_AMT22_flow_cytometry | prochlorococcus_abundance_P701A90Z_Tarran | https://simonscmap.com/catalog/datasets/JC079_AMT22_flow_cytometry |
| 42 | tblJC079_AMT22_flow_cytometry | synechococcus_abundance_P700A90Z_Tarran | https://simonscmap.com/catalog/datasets/JC079_AMT22_flow_cytometry |
| 43 | tblJC079_AMT22_flow_cytometry | picoeukaryotic_abundance_PYEUA00A_Tarran | https://simonscmap.com/catalog/datasets/JC079_AMT22_flow_cytometry |
| 44 | tblJR20131005_AMT23_flow_cytometry | prochlorococcus_abundance_P701A90Z_Tarran | https://simonscmap.com/catalog/datasets/JR20131005_AMT23_flow_cytometry |
| 45 | tblJR20131005_AMT23_flow_cytometry | synechococcus_abundance_P700A90Z_Tarran | https://simonscmap.com/catalog/datasets/JR20131005_AMT23_flow_cytometry |
| 46 | tblJR20131005_AMT23_flow_cytometry | picoeukaryotic_abundance_PYEUA00A_Tarran | https://simonscmap.com/catalog/datasets/JR20131005_AMT23_flow_cytometry |
| 47 | tblJR20140922_AMT24_flow_cytometry | prochlorococcus_abundance_P701A90Z_Tarran | https://simonscmap.com/catalog/datasets/JR20140922_AMT24_flow_cytometry |
| 48 | tblJR20140922_AMT24_flow_cytometry | synechococcus_abundance_P700A90Z_Tarran | https://simonscmap.com/catalog/datasets/JR20140922_AMT24_flow_cytometry |
| 49 | tblJR20140922_AMT24_flow_cytometry | picoeukaryotic_abundance_PYEUA00A_Tarran | https://simonscmap.com/catalog/datasets/JR20140922_AMT24_flow_cytometry |
| 50 | tblJR15001_AMT25_flow_cytometry | prochlorococcus_abundance_P701A90Z_Tarran | https://simonscmap.com/catalog/datasets/JR15001_AMT25_flow_cytometry |
| 51 | tblJR15001_AMT25_flow_cytometry | synechococcus_abundance_P700A90Z_Tarran | https://simonscmap.com/catalog/datasets/JR15001_AMT25_flow_cytometry |
| 52 | tblJR15001_AMT25_flow_cytometry | picoeukaryotic_abundance_PYEUA00A_Tarran | https://simonscmap.com/catalog/datasets/JR15001_AMT25_flow_cytometry |
| 53 | tblDY110_AMT29_flow_cytometry | prochlorococcus_abundance_P701A90Z_Tarran | https://simonscmap.com/catalog/datasets/DY110_AMT29_flow_cytometry |
| 54 | tblDY110_AMT29_flow_cytometry | synechococcus_abundance_P700A90Z_Tarran | https://simonscmap.com/catalog/datasets/DY110_AMT29_flow_cytometry |
| 55 | tblDY110_AMT29_flow_cytometry | picoeukaryotic_abundance_PYEUA00A_Tarran | https://simonscmap.com/catalog/datasets/DY110_AMT29_flow_cytometry |

Table S2: List of environmental variables colocalized with the observations of cyanobacteria.

| | Variable | Unit | Long Name | Acquisition Method | Start | End | Data Source | Distributor |
|---|---|---|---|---|---|---|---|---|
| 1 | SST | $^{o}$C | Sea Surface Temperature | Satellite | 1981-09-01 | 2019-04-27 | National Centers for Environmental Information | https://podaac.jpl.nasa.gov/ |
| 2 | CHL | mg/m$^3$ | Chlorophyll concentration - 8day averaged | Satellite | 1998-01-01 | 2018-06-26 | Copernicus-GlobColour | http://marine.copernicus.eu/ |
| 3 | SSS | psu | Sea Surface Salinity | Satellite | 2015-03-31 | 2019-04-21 | Remote Sensing Systems, Santa Rosa, CA, USA http://www.remss.com/missions/smap/ | https://podaac-opendap.jpl.nasa.gov/opendap/allData/smap/L3/RSS/V3/8day_running/SCI/70KM |
| 4 | PAR | einstein m$^{-2}$ day$^{-1}$ | Photosynthetically Available Radiation | Satellite | 2015-01-01 | 2020-01-15 | Ocean Biology Processing Group at NASA GSFC | Ocean Biology Processing Group at NASA GSFC |
| 5 | SLA | m | Sea Level Anomaly | Satellite | 1993-01-01 | 2018-06-10 | SL-TAC | http://marine.copernicus.eu/ |
| 6 | ADT | m | Absolute Dynamic Topography | Satellite | 1993-01-01 | 2018-06-10 | SL-TAC | http://marine.copernicus.eu/ |
| 7 | Ugosa | m/s | Geostrophic velocity anomalies: zonal component | Satellite | 1993-01-01 | 2018-06-10 | SL-TAC | http://marine.copernicus.eu/ |
| 8 | Vgosa | m/s | Geostrophic velocity anomalies: meridian component | Satellite | 1993-01-01 | 2018-06-10 | SL-TAC | http://marine.copernicus.eu/ |
| 9 | $NO_3$ | mmol/m$^3$ | Mole concentration of Nitrate in sea water | Blend | 2011-12-31 | 2019-04-27 | MERCATOR BIOMER4V1R2 | http://marine.copernicus.eu |
| 10 | $PO_4$ | mmol/m$^3$ | Mole concentration of Phosphate in sea water | Blend | 2011-12-31 | 2019-04-27 | MERCATOR BIOMER4V1R2 | http://marine.copernicus.eu |
| 11 | Fe | mmol/m$^3$ | Mole concentration of dissolved iron in sea water | Blend | 2011-12-31 | 2019-04-27 | MERCATOR BIOMER4V1R2 | http://marine.copernicus.eu |
| 12 | $O_2$ | mmol/m$^3$ | Mole Concentration of dissolved Oxygen in sea water | Blend | 2011-12-31 | 2019-04-27 | MERCATOR BIOMER4V1R2 | http://marine.copernicus.eu |
| 13 | Si | umol/L | Mole concentration of Silicate in sea water | Blend | 2011-12-31 | 2019-04-27 | MERCATOR BIOMER4V1R2 | http://marine.copernicus.eu |
| 14 | PP | g/m$^3$/day | Net primary productivity of Carbon per unit volume | Blend | 2011-12-31 | 2019-04-27 | MERCATOR BIOMER4V1R2 | http://marine.copernicus.eu |
| 15 | Density_WOA_clim | kg/m$^3$ | Objectively Analyzed Climatology for Density World Ocean Atlas | In-Situ | Monthly Climatology | Monthly Climatology | http://doi.org/10.7289/V5NZ85MT | https://www.nodc.noaa.gov/OC5/woa13/ |
| 16 | Salinity_WOA_clim | psu | Objectively Analyzed Climatology for Salinity World Ocean Atlas | In-Situ | Monthly Climatology | Monthly Climatology | http://doi.org/10.7289/V5NZ85MT | https://www.nodc.noaa.gov/OC5/woa13/ |
| 17 | Nitrate_WOA_clim | mcmol/L | Objectively Analyzed Climatology for Nitrate World Ocean Atlas | In-Situ | Monthly Climatology | Monthly Climatology | http://doi.org/10.7289/V5NZ85MT | https://www.nodc.noaa.gov/OC5/woa13/ |
| 18 | Phosphate_WOA_clim | mcmol/L | Objectively Analyzed Climatology for Phosphate World Ocean Atlas | In-Situ | Monthly Climatology | Monthly Climatology | http://doi.org/10.7289/V5NZ85MT | https://www.nodc.noaa.gov/OC5/woa13/ |
| 19 | Silicate_WOA_clim | mcmol/L | Objectively Analyzed Climatology for Silicate World Ocean Atlas | In-Situ | Monthly Climatology | Monthly Climatology | http://doi.org/10.7289/V5NZ85MT | https://www.nodc.noaa.gov/OC5/woa13/ |
| 20 | Oxygen_WOA_clim | mcmol/L | Objectively Analyzed Climatology for Dissolved Oxygen World Ocean Atlas | In-Situ | Monthly Climatology | Monthly Climatology | http://doi.org/10.7289/V5NZ85MT | https://www.nodc.noaa.gov/OC5/woa13/ |
